# Functional and structural consequences of epithelial cell invasion by *Bordetella pertussis* adenylate cyclase toxin

**DOI:** 10.1101/2020.01.31.928192

**Authors:** Christelle Angely, Daniel Ladant, Emmanuelle Planus, Bruno Louis, Marcel Filoche, Alexandre Chenal, Daniel Isabey

## Abstract

*Bordetella pertussis*, the causative agent of whopping cough, produces an adenylate cyclase toxin (CyaA) that plays a key role in the host colonization by targeting innate immune cells which express CD11b/CD18, the cellular receptor of CyaA. CyaA is also able to invade non-phagocytic cells, via a unique entry pathway consisting in a direct translocation of its catalytic domain across the cytoplasmic membrane of the cells. Within the cells, CyaA is activated by calmodulin to produce high levels of cyclic adenosine monophosphate (cAMP) and alter cellular physiology. In this study, we explored the effects of CyaA toxin on the cellular and molecular structure remodeling of A549 alveolar epithelial cells. Using classical imaging techniques, biochemical and functional tests, as well as advanced cell mechanics method, we quantify the structural and functional consequences of the massive increase of intracellular cyclic AMP induced by the toxin: cell shape rounding associated to adhesion weakening process, actin structure remodeling for the cortical and dense components, increase in cytoskeleton stiffness, and inhibition of migration and repair. We also show that, at the low concentrations that may be found *in vivo* during *B. pertussis* infection, CyaA impairs the migration and wound healing capacities of the intoxicated alveolar epithelial cells. Our results suggest that the CyaA, beyond its major role in disabling innate immune cells, might also contribute to the local alteration of the epithelial barrier of the respiratory tract, that is an hallmark of *pertussis*.

## Introduction

The adenylate cyclase (CyaA) is a major toxin secreted by *Bordetella pertussis*, the causative agent of whopping cough. This toxin plays a key role in the early stage of colonization of the respiratory tract by *Bordetella pertussis*. CyaA is able to invade eukaryotic cells in which it translocates its catalytic domain which is activated by endogenous calmodulin to catalyze a massive production of cyclic AMP (cAMP), resulting in profound alterations of cellular physiology [1–3]. The cytotoxic effects of CyaA are mainly directed towards neutrophils and macrophages, as well as other innate immune cells that express the CD11b/CD18 integrin, which is the main cellular receptor of CyaA [4]. CyaA inhibits the phagocytic function of neutrophils and macrophages by impairing chemotaxis and oxidative response, and eventually triggers cell apoptosis and necrosis [5–9]. Yet, CyaA is also able to invade a wide variety of non-phagocytic cells [1, 2, 8]. A similar unique mechanism of entry in the cytosol is likely used by CyaA in both immune and non-immune cells. It consists in a direct translocation of the catalytic domain across the cytoplasmic membrane of the cells [10] which is confirmed by the rapidity of the intoxication (or internalization) process (basically a few seconds) with however differences depending on cell type [10–12]. Nevertheless, the molecular mechanisms by which CyaA penetrates the target cells are still largely elusive and the physiological consequences of the toxin activity remain to be precisely determined especially in non immune cells [1]. In particular, several studies have recently suggested that CyaA might also act at the site of infection on the epithelial cells of the respiratory tract. First, Eby *et al.* [13] have quantified the amounts of CyaA that is produced locally during infection of the respiratory tract and they found that, at the bacterium-target cell interface, the concentration of CyaA may exceed 100 ng/ml (about 0.6 nM of toxin). Other studies have suggested that CyaA at these concentrations may affect the epithelial cells and eventually contribute to the alteration of the epithelial barrier which is primarily affected by the cytopathic action of the tracheal cytotoxin, TCT, released by *B. pertussis*. Ohnishi *et al.* (2008) first showed that CyaA induces morphological changes on cultured rat alveolar epithelial cells [14]. Eby *et al.* reported that polarized T84 cell monolayers as well as human airway epithelial cultures could respond to nanomolar concentrations of CyaA when it was applied to the basolateral membranes [15]. We then showed that CyaA can invade A549 alveolar epithelial cells and trigger a significant remodeling of their molecular adhesion systems [16] and, more recently, Hasan *et al.* (2018) analyzed the impact of CyaA on the functional integrity of human bronchial epithelial cells cultured at the air-liquid interface [17].

Cytoskeletal (CSK) rearrangement may be caused by different signal events including modifications in the intracellular Ca^2+^ and cAMP-mediated signaling which have been involved in perturbations of the actin CSK homeostasis in different cells [18, 19]. These two factors are indeed deeply modified and markedly increased after CyaA invasion suggesting that the CyaA toxin could trigger significant CSK remodeling of the host cells [8, 9, 20, 21].

In the present study, we further characterized the CyaA-induced structural remodeling and functional alterations of the CSK and mechanical properties of A549 alveolar epithelial cells. We show that CyaA intoxication reduces the migration and wound healing capacities of these cells when they are exposed to toxin doses mimicking those reached in vivo during *B. pertussis* infection. Our present results therefore suggest that the CyaA toxin may contribute to the local disruption of the integrity of the airway epithelium.

## Materials and methods

### Cellular Model of Intoxication

#### Culture of Alveolar Epithelial Cell lines (AECs)

Experiments were carried out on A549 cells which are an alveolar epithelial cell line (AECs) classically used for cell respiratory physiology studies. Briefly, this line, which originates from a pulmonary epithelium adenocarcinoma taken from patient, is obtained from the National Cancer Institute’s lineage library (ref: ATCC Collection No. CCL-185). It is a hypo-triploid line that has 24% of 12-chromosome cells, 22% of 64-chromosome cells and the rest of diploid cells. In addition, A549-type epithelial cells have been used in the laboratory for many years [22, 23] as they express a phenotype like certain pulmonary alveolar epithelial cells, i.e., the type II pneumocytes [24].

AECs offer many advantages for studying in vitro the pathophysiological response of pulmonary cells [25]. They form adherent and tight junctions when grown to confluence and express a wide variety of cytokines, growth factor and receptors and notably several transmembrane receptors of the integrin type: β_1_; β_3_; α_3_; α_6_; α_5_ and α_2_ [26]. These integrin receptors bind the synthetic peptide containing the RGD sequence present in many extracellular matrix components. The peptide RGD is classically used for integrin-specific cell-binding as done in the present study and in many previous studies [27, 28]. To maintain integrin expression at a sufficiently high level [29], the passage number was maintained in the low range (≈ 12^th^ 16^th^).

The cells are cultured in plastic flasks treated for cell adhesion with a filter cap (25 or 75 cm^2^, Techno Plastic Products AG, Switzerland). The culture medium consists of DMEM (Gibco Life Technologies), 10% fetal calf serum or FCS (Sigma-Aldrich, St. Louis, MO, USA) as well as 1% antibiotics (penicillin and streptomycin). The FCS is the most complex component because it contains growth factors, hormones, elements of the extracellular matrix, e.g., fibronectin and vitronectin, and all other element contained in the blood, except the figured elements, i.e., the coagulation factors and the complement.

The cultures are incubated at 37 °C in a controlled atmosphere (5% CO_2_ and 95% humidity). Cells consume nutrients from the environment and produce metabolites. It is therefore necessary to regularly renew the medium, namely, every 24 or 48 hours. The cells are adherent to the support and must therefore be peeled off using trypsin-EDTA 0.05% (Sigma-Aldrich, St. Louis, MO, USA) and then subcultured with a split ratio of 1/10. After centrifugation at 200 g, the cell pellet is re-suspended in DMEM-10% FCS medium and a part is transferred to another flask. To keep the line and have a stock, the unused cells are frozen. For experiments, we used a density of 7.10^5^ cells and seeded on petri-dish (TPP *ϕ*34mm) coated with fibronectin at 10 ng/mm^2^.

#### Production and Purification of CyaA and its Inactive Variant CyaAE5

CyaA and the enzymatically inactive variant CyaAE5 (resulting from a LQ dipeptide insertion between D188 and I189 in the catalytic core of the enzyme [30]) were expressed in E. coli and purified to homogeneity by previously established procedures [4, 31]. Succinctly, the inclusion bodies were solubilized overnight at 4 °C in 20 mM Hepes, 8 M urea, pH 7.4. The soluble urea fraction was recovered by centrifugation, supplemented with 0.14 M NaCl and then loaded on a Q-Sepharose fast flow resin equilibrated with 20 mM Hepes, 140 NaCl, 8 M urea, pH 7.4. Contaminants were eliminated by an extensive wash in the same buffer; the CyaA protein is then eluted using a NaCl gradient (CyaA elution occurs around 500 mM NaCl). After dilution of the eluate in 20 mM Hepes, 8 M urea to reach 100 mM NaCl, the CyaA batch is loaded onto a second Q-sepharose Hi Performance column. Washing and elution are operated in the same conditions as the first Q. This step allows to obtain a concentrated CyaA protein which is then diluted 5 times with 20 mM Hepes, 1 M NaCl, pH 7.4 and loads onto a 70 ml phenyl-sepharose column and washes with 20 mM Hepes, 1 M NaCl, with Hepes 20 mM again and then with 50% isopropanol. After an extensive wash, the toxin is eluated with 20 mM Hepes and 8 M urea. The eluate is then applied onto a sephacryl 500 (GE healthcare, HIPREP 26/60) equilibrated in 20 mM Hepes and 8 M urea. CyaA batches are pooled and concentrated by ultrafiltration and stored at 20 °C in 20 mM Hepes and 8 M urea. CyaA toxin concentration is determined by UV-spectrophotometry using a molecular extinction coefficient Em280 = 143590 M^−1^cm^−1^ computed from the CyaA sequence on the ProtParam server (http://web.expasy.org/protparam/). The purity of CyaA batches is higher than 90% as judged by SDS PAGE analysis and contained less than 1 EU of LPS/μg of protein as determined by a standard LAL assay (Lonza). Finally, CyaA is refolded into a urea-free, monomeric and functional holo-state [31, 32]. The refolding efficiency is around 40 ± 5% (population monomer / total population of proteins). The biological functions, i.e., hemolysis, translocation of ACD and cAMP production, are routinely assayed as described in [31] and [33]. The aliquots of monomeric species of CyaA are stored at −20 °C in 20 mM Hepes, 150 mM NaCl, 2 mM CaCl_2_, pH 7.4.

#### Handling of CyaA and CyaAE5 Toxins

For our experiments, CyaA and CyaAE5 recombinant proteins are diluted into a mix solution containing DMEM-10% SVF, buffer and CaCl2. CyaA mixture or CyaAE5 mixture is directly added on cells for 30 or 60 min. We used a range of concentration of CyaA or CaAE5 toxin from 0.5 to 10 nM.

### Biological Tests

#### cAMP Assays

A549 cells are seeded at 3.10^5^ cells/well in 96 well plates. After 24 hours, cells are exposed for 60 min to three different concentrations (0.5, 5, and 10 nM) of either the CyaA toxin or the enzymatically inactive CyaAE5 protein. The CyaA mix was removed and cells were washed. Cells were then recovered and lysed using 0.1 M HCl in order to collect the intracellular content. The concentration of cAMP in the lysates was then measured by using a competitive ELISA kit for cAMP (Invitrogen, ref. EMSCAMPL) following the recommendations of the manufacturer.

#### Viability Evaluation by MTT Test

MTT (3- [4,5-dimethyl-2-thiazolyl] −2,5-diphenyltetrazolium bromide, Sigma M5655) is a tetrazolium salt giving a yellow solution when diluted in the medium. It is converted into insoluble violet formazan in the culture medium after cleavage of the tetrazolium ring by the active mitochondrial dehydrogenases of living cells only (dead cells are not detected by this assay). 2500 cells are seeded per well in a 96-well plate. After 24 hours of incubation, the cells are synchronized with DMEM at 0% FCS without phenol red. After 72 hours of incubation, different concentrations of CyaA are added for 60 minutes. Then the CyaA mix was removed and cells were washed. Finally, 50 μL of MTT solution at 2 μg/ml are then added to each well. After 4 hours of incubation, the medium is removed by inversion. 200 μL of pure DMSO were then added to each well. Finally, the 96-well plate is read from the ELISA reader at a wavelength of 550 nm. Results are expressed in terms of optical density OD (or absorbance).

#### Actin Cytoskeleton and Focal Adhesion Staining

The A549 cells are seeded at a density of 75,000 cells per coverslip (12 mm). After 24h incubation at 37 °C and 5% CO_2_, the cells reach 60% confluence and are incubated for 60 min at 37 °C in complete medium (control conditions) or complete medium supplemented with CyaA toxin (0.5, 5 or 10 nM final concentrations). In some experiments, in control and after 60 min of CyaA incubation, cells are exposed 30 min to Mn^2+^ (0.5 mM final concentration). The cells are washed with PBS and then fixed for 10 min in 4% paraformaldehyde, permeabilized by addition of 0.3% Triton in PBS. Non-specific sites are blocked by incubation of PBS-1% BSA during 30 min. Then, cells are incubated for 1 hour at room temperature with anti-phosphotyrosine primary antibody PY99 (sc-7020, Santa Cruz Biotechnology) diluted at 1/150 in PBS-1% BSA. The PY99 antibody is then detected with an Alexa-488 labeled secondary antibody (Life Technology ref. A-21042 diluted 1/1000 in PBS) to reveal focal adhesion points (in green). After rinsing with PBS-0.02% Triton, the cells are incubated in a dilute solution of phalloidin tetramethylrhodamine bisothiocynate (Sigma, ref. P1951 diluted at 1/2000 in PBS) in order to display actin fibers (in red).

Coverslips were mounted on a slide with the cell side down in Prolong (LifeTechnology, ref. P36974). Staining structures are observed, and images are acquired on an Axio Imager confocal microscope (Zeiss) at 63× magnification. To analyze cellular images, the mean level of fluorescence of F-actin is determined in each cell by automatic quantification by ImageJ software relatively to the fluorescent background level of each image. The effective mean fluorescence of cells is obtained after subtraction of background fluorescence. In order to distinguish the cortical and the dense actin filaments, two different fluorescence thresholds on ImageJ software were used, i.e., 152 for the cortical actin and 534 for the dense actin, to permit the quantification of actin fluorescence at two detected levels. In the same way, different thresholds were used to distinguish different size areas of focal adhesion points (< or > to 1 μm^2^). Finally, on the same coverslips, we used the LSM5 Pascal software to make stacks of images (every 0.5 μm) in the vertical direction ‘z’ and to carry out a 3D reconstruction of the shape of the cell in order to measure the cell height in each condition.

### Functional assessments

#### Measure of CSK stiffness by MTC

A549 cell mechanical properties were measured by a bead micromanipulation system through CSK specific probing, called Magnetic Twisting Cytometry (MTC) which has been extensively described in the literature [27, 28]. Probes are ferromagnetic microbeads coated with an integrin specific ligand, e.g., fibronectin. When an external magnetic field is exerted, these beads apply a torque to the living cell structure through CSK specific-ligand binding. The relationship between the measured bead rotation angle (angular strain averaged over the cell culture during cellular loading), and the applied torque provides CSK-specific measurements of mechanical (rheological) properties of the cell. This technique invented by Wang *et al.* in 1993 [28] has been recently upgraded by Isabey and co-workers to achieve a multiscale quantification of cellular and molecular parameters [27]. Each MTC measurement is performed twice on a wide number of cells (7 × 10^4^). The large number of beads and their uniform distribution throughout the cell culture guarantee an instantaneous homogenized cell response, representative of the dominant structural behavior. The key parameter obtained is the elasticity modulus or cytoskeleton stiffness (*E* in Pa), which has been measured for the different concentrations of the CyaA toxin. *E* quantifies the mechanical stiffness of CSK structure and reflects also the intracellular tension or prestress [34].

#### Measure of Cell Migration and Wound Healing

A549 cells were seeded at a density of 10^5^ cells/chamber in a 12-well simple chamber. There were incubated during 24 hours in complete medium (DMEM, 1% penicillin, 5% SVF) to form a confluent monolayer. Cells were exposed to the different concentrations of CyaA toxin or to complete medium (control conditions). The cell monolayer was scratched with a pipette tip of 5 μm-distal diameter to create a wound and rinsed 3 times with medium in order to remove the entire detached cell population. Transmission images of the cell monolayer with the wound are taken every 3 hours with a black and white CDD camera (Kappa DX20-H) mounted on a fluorescent-transmission microscope (Leitz Labovert FS). Cell migration was determined by the aptitude of cells to close the wound. The quantification of the repair of the monolayer is made by measuring the wound area using ImageJ (W.S. Rasband; NIH, Bethesda). The evolution of the repaired area is quantified by the following formula:

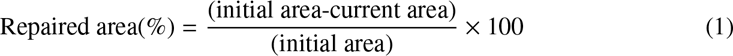

The wound repair is followed until complete closure (repaired area=100%).

## Results

We previously showed [16] that A549 alveolar epithelial cells are sensitive to CyaA that triggers a cAMP-dependent remodeling of their molecular adhesion system, resulting in a significant weakening in their initial adhesion processes. Here we further examined the effects of CyaA on the reorganization of the actin cytoskeleton and evolution of focal adhesions. A549 cells were exposed for 1 hr to various concentrations of CyaA (ranging form 0.5 to 10 nM) and then the F-actin structures were visualized by phalloidin staining (in red) and focal adhesions were visualized with anti-phosphotyrosin PY99 antibodies (in green) (Fig 1).

**Fig 1.**
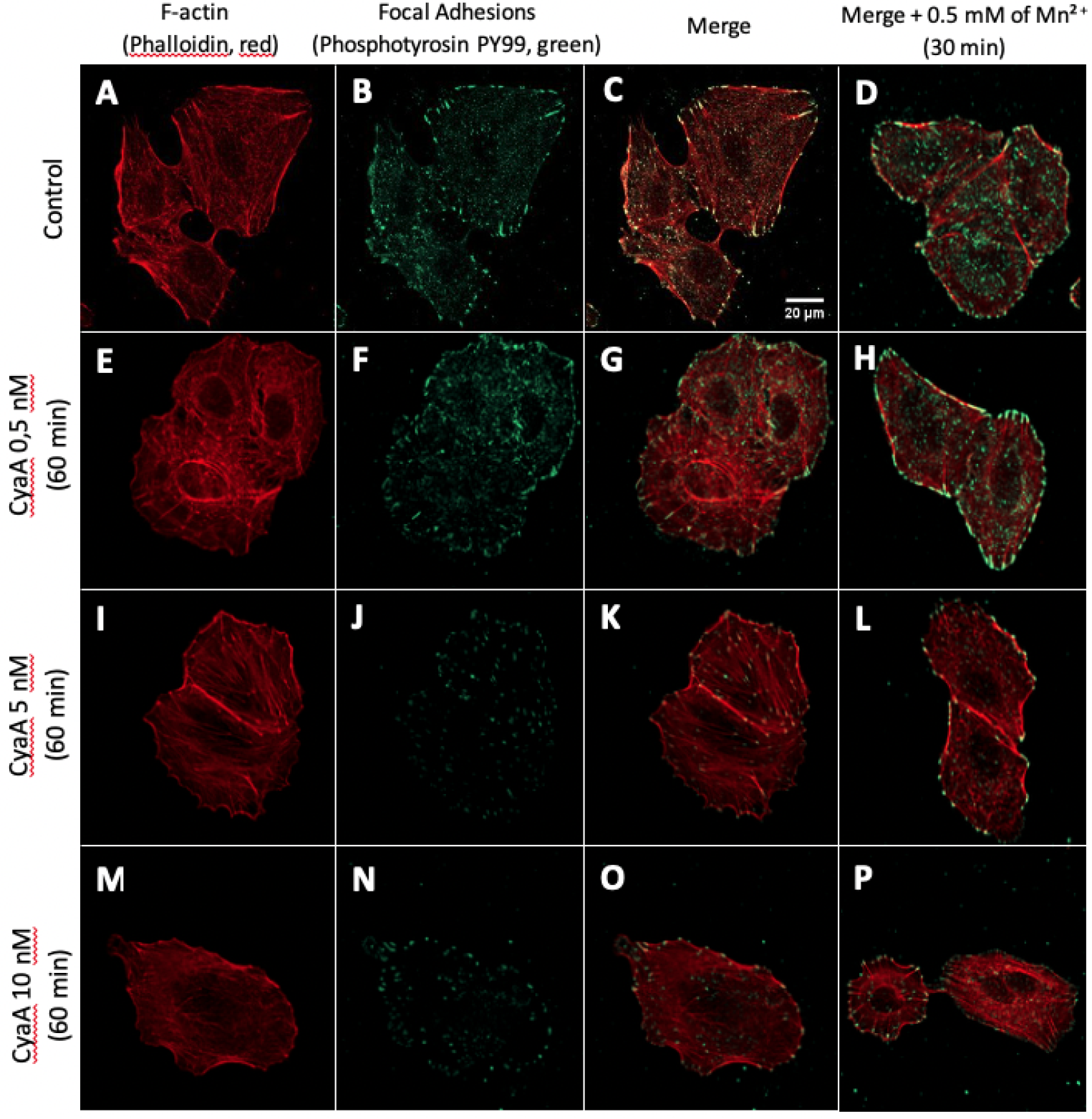
Cell imaging of F-actin structure and focal adhesion points. Co-staining of F-actin (with phalloidin, in red) and of focal adhesions (with anti-phosphotyrosin PY99 antibody, in green) in fixed A549 cells (control conditions) after 1 hour of exposure to the indicated concentrations of CyaA toxin. (A, E, I, M) F-actin staining, (B, F, J, N) focal adhesion staining, (C, G, K, O) merge images. Right panels (D, H, L, P) show merged images of F-actin staining and focal adhesion staining for cells incubated with the indicated concentrations of CyaA for 60 min followed by a 30 min exposure to Mn^2+^ at a concentration of 0.5 mM. (A-D) Control conditions (no CyaA), (E-H) cell exposure to 0.5 nM of CyaA, (I-L) cell exposure to 5 nM of CyaA, (M-P) cell exposure to 10 nM of CyaA. Images were obtained by confocal microscopy with ×63 magnification. Fig S1 and Fig S2 further document the cell viability and intracellular cAMP accumulation in the A549 cells exposed to CyaA.

In cells incubated with CyaA, a marked remodeling of actin structure and focal adhesion reorganization is observed. This effect is characterized by a progressive concentration-dependent disappearance of stress fibers associated with a significant loss of number of focal adhesion points. Noteworthy, the effect of CyaA intoxication on both actin structure and focal adhesion could be partially counteracted by addition of manganese (0.5 mM Mn^2+^) during the last 30 min of incubation with CyaA (Fig 1). Control cells exposed to Mn^2+^ (Fig 1D) shows a strong reinforcement of both F-actin structure and most particularly focal adhesion points, as expected given the activation of integrins forced by Mn^2+^ [35, 36]. It appears that CyaA-intoxicated A549 cells still respond to manganese exposure but to a lesser extent than the non-intoxicated (control) cells.

Quantifications of global fluorescent F-actin at the basal face of spread A549 after 60 min of CyaA exposure (Fig 2) reveal a significant decrease in F-actin intensity, in a toxin concentration-dependent manner. This confirms the microscopic observations shown above. Quantifications of F-actin structures at low density F-actin (cortical or sub-membranous F-actin) and high-density F-actin (stress fibers) reveal that cortical actin sub-structures are predominantly and significantly affected by CyaA exposure while the dense actin sub-structures are only slightly affected (Fig 2). The decrease in total F-actin intensity in CyaA-intoxicated cells is also observed after Mn^2+^ exposure although the distribution of cortical versus dense F-actin is modified in favor of the dense F-actin with manganese (compare Fig 2 and Fig 3).

**Fig 2.**
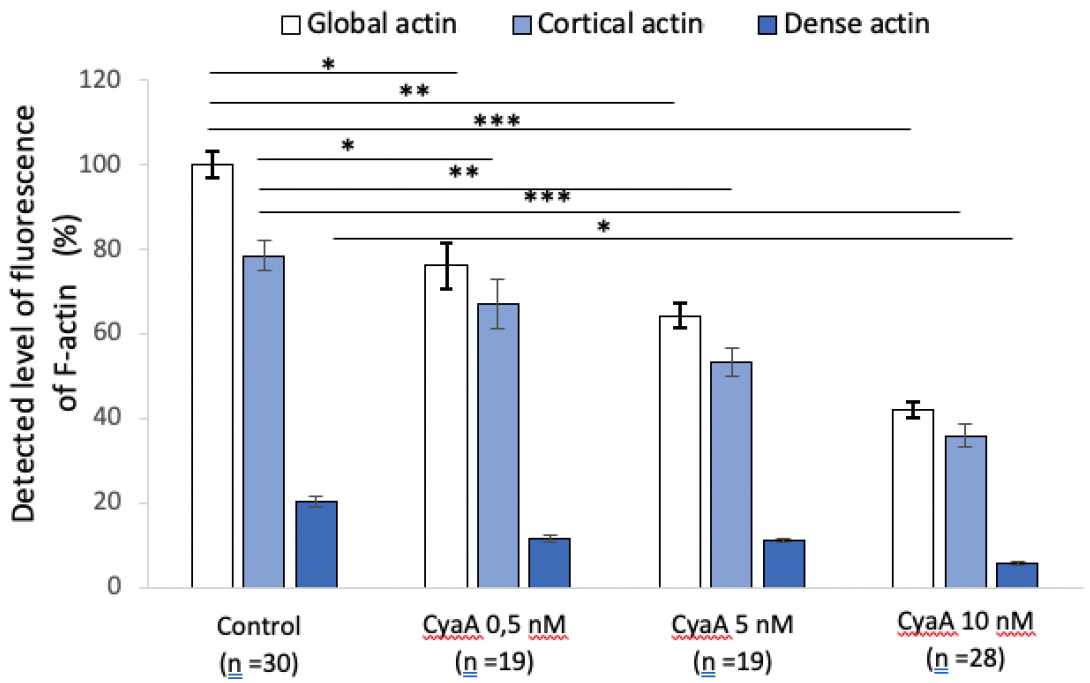
Quantification of fluorescence levels of global, cortical and dense F-actin in A549 after 60 min of CyaA exposure. The bar graph provides a quantification of F-actin fluorescence in A549 cells before (control conditions) and after 60 min of exposure to the indicated concentrations of CyaA. The specific distribution of the cortical (submembranous) and dense (stress fibers) actin structures were analyzed by using intermediate thresholds of fluorescence (see Material and Methods). ‘n’ corresponds to the number of analyzed cells. Error bars are ± SEM; * *p* ≤ 0.05; ** *p* ≤ 0.01; *** *p* ≤ 0.001.

**Fig 3.**
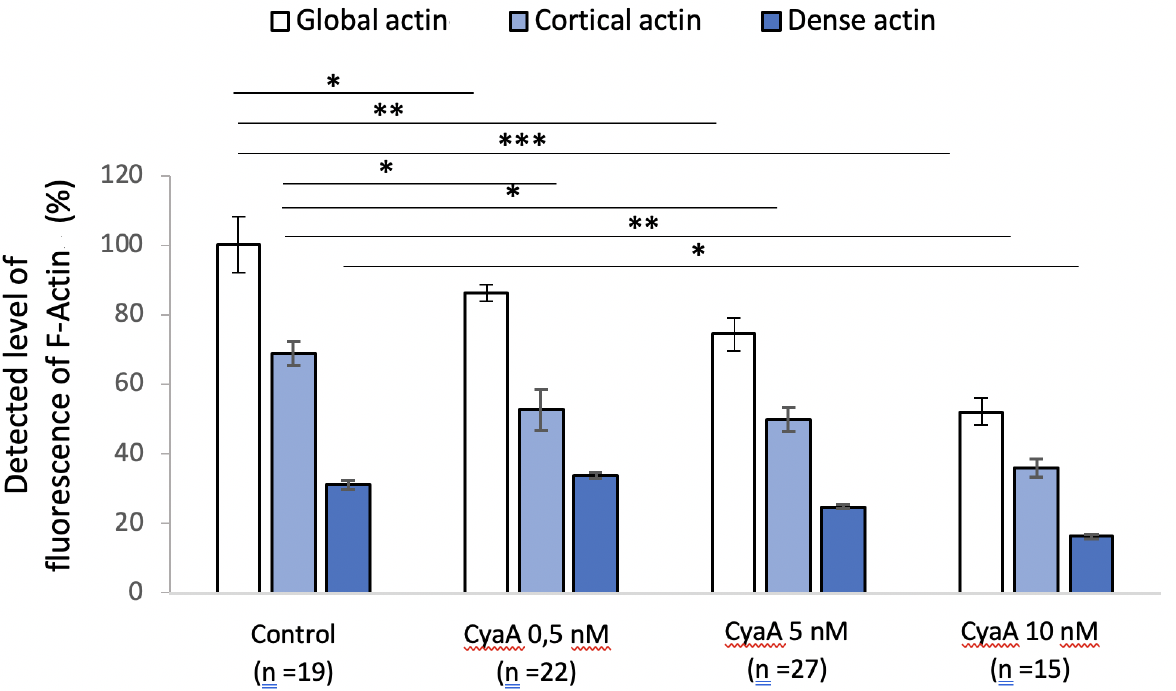
Quantification of fluorescence levels of global, cortical and dense F-actin in A549 cells after 60 min of CyaA followed by 30 min of Mn^2+^ exposure. The bar graph provides a quantification of F-actin fluorescence in A549 cells before (control conditions) and after 60 min exposure to the indicated concentrations of CyaA, then, after washing, a 30 min exposure to Mn^2+^ at 0.5 mM. The specific distribution of the cortical (submembranous) and dense (stress fibers) actin structures were analyzed by using intermediate thresholds of fluorescence (see Material and Methods). ‘n’ corresponds to the number of analyzed cells. Error bars are ± SEM; * *p* ≤ 0.05; ** *p* ≤ 0.01; *** *p* ≤ 0.001.

Quantification of the number of focal adhesion (FA) points after 60 min of CyaA exposure is given for all FA sizes as well as for the small (≤ 1 μm^2^) and large (> 1 μm^2^) FA sizes (Fig 4). The total number of adhesion sites in cells exposed to CyaA markedly decreases as the toxin concentration increases. For all conditions studied, the wide majority of adhesion sites is constituted by small adhesion sites, the number of which is significantly reduced as the CyaA concentration increases. By contrast, the number of large adhesion sites is always significantly smaller than the total number of FA (statistics not presented) but, compared to small FA, the decay in the number of large FA (as the CyaA concentration increases) is only slightly significant (Fig 4). The exposure to Mn^2+^ (Fig 5) modifies the size distribution of FA by decreasing the proportion of small adhesion sites while increasing that of large adhesion sites (Fig 5). Thus, CyaA triggers a drastic weakening in cell adhesion processes which can be partly reversible by exposure of cells to Mn^2+^. In other words, manganese treatment partially counterbalances the changes in FA size and remodeling of F-actin structures induced by the toxin.

**Fig 4.**
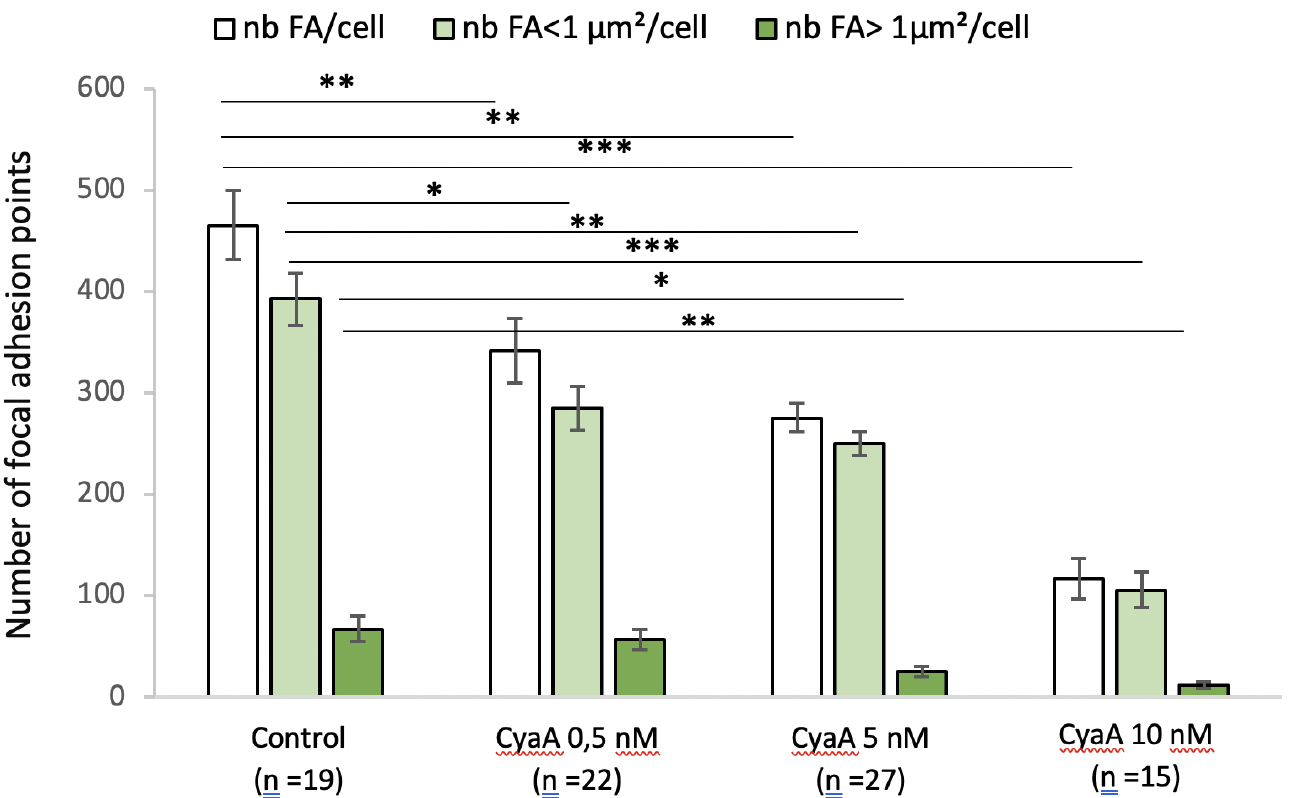
Quantification of the number and size of focal adhesion points of A549 cells exposed to CyaA. This bar graph shows the numbers of focal adhesion (FA) point per cell and their distribution in terms of size (surface areas of FA below and above 1 μm^2^) in A549 cells incubated for 60 min with (or without control conditions) the indicated concentrations of CyaA. ‘n’ corresponds to the number of analyzed cells. Error bars are ± SEM; * *p* ≤ 0.05; ** *p* ≤ 0.01; *** *p* ≤ 0.001.

**Fig 5.**
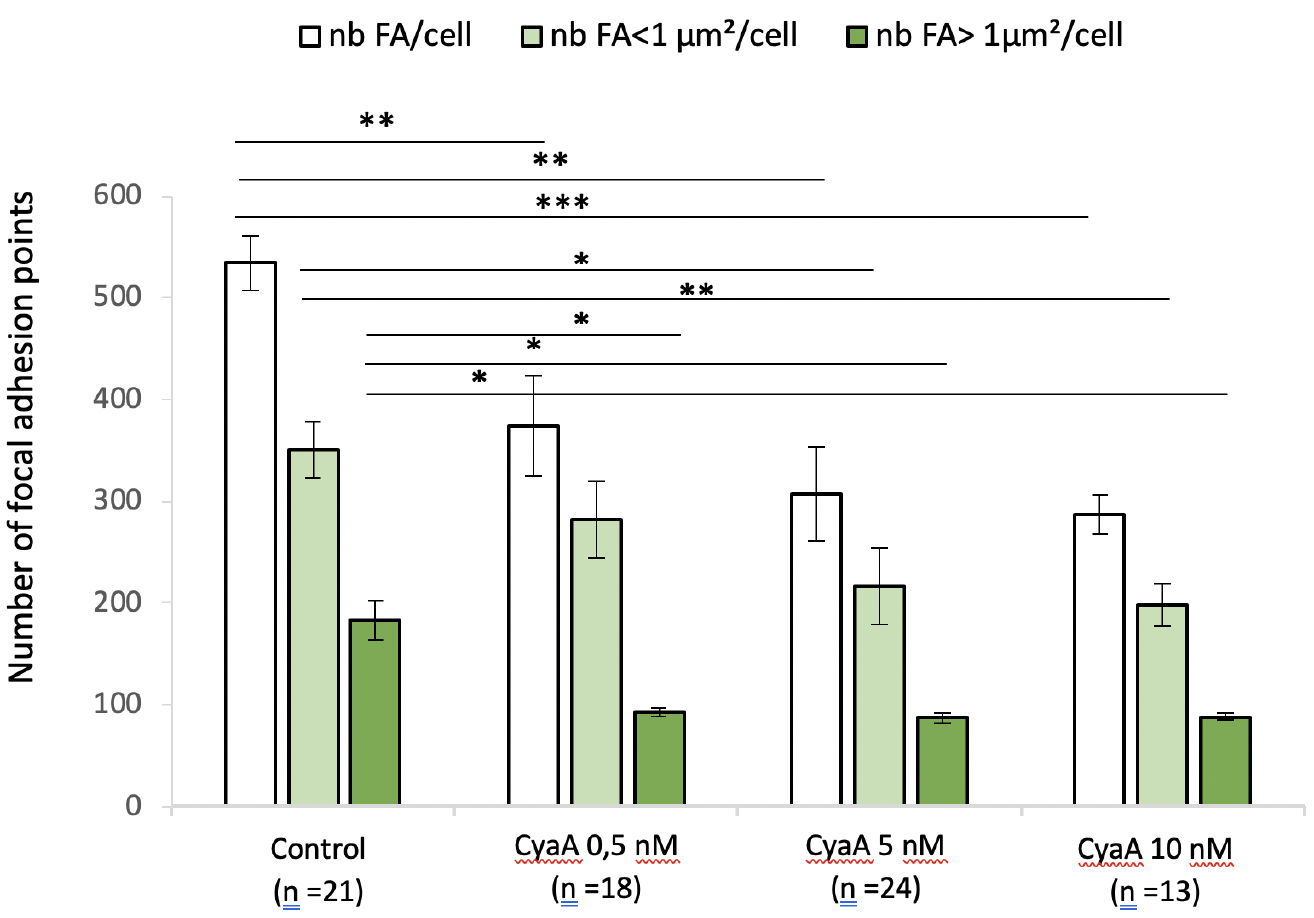
Quantification of the number and size of focal adhesion points of A549 cells exposed to CyaA followed by 30 min of Mn^2+^ exposure. This bar graph shows the numbers of focal adhesion (FA) point per cell and their distribution in terms of size (surface areas of FA below and above 1 μm^2^) in A549 cells before (control conditions) and after 60 min exposure to the indicated concentrations of CyaA, then, after washing, a 30 min exposure to Mn^2+^ at 0.5 mM. ‘n’ corresponds to the number of analyzed cells. Error bars are ± SEM; * *p* ≤ 0.05; ** *p* ≤ 0.01; *** *p* ≤ 0.001.

The effects of CyaA on the global shape of A549 cells were examined by analyzing images from different confocal planes. Statistics of cell spreading area and cell height for the different conditions of CyaA exposure tested reveals significant changes in cell shape after intoxication by the CyaA toxin (Fig 6A and 6B). The cell shape change can logically be related to the CyaA induced weakening process of adhesion presented in Fig 1 to 5 and suggests that CyaA triggers cell rounding. This is consistent with prior studies that showed a similar change in cell morphology, i.e., namely a rounding process, in various cultured cell lines - notably in type II alveolar cells [14].

**Fig 6.**
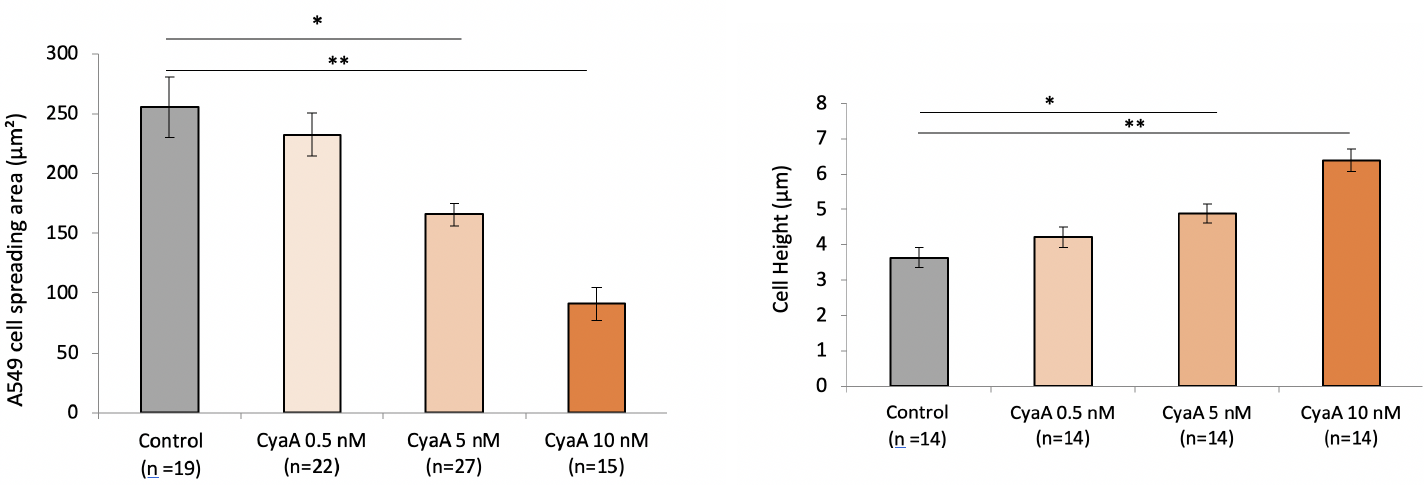
Effect of CyaA on A549 cell rounding. A549 cells were exposed for 60 min to the indicated concentrations of CyaA toxin (control=no CyaA) and analyzed by confocal microscopy to determine (A) the cell spreading area (in μm^2^) which is measured in the basal plane of the *z*-stack confocal images and (B) cell height (in μm) which is measured from the distance separating the basal and apical planes of the *z*-stack confocal images. ‘n’ corresponds to the number of analyzed cells. Differences between the different groups are quantified by ANOVA test compared to the control conditions. Error bars are ± SEM; * *p* ≤ 0.05; ** *p* ≤ 0.01; *** *p* ≤ 0.001.

To examine whether CyaA intoxication affects acto-myosin motors and notably intracellular cytoskeletal tension, we measured the changes in the cytoskeleton (CSK) stiffness. We analyzed the CSK stiffness by using the Magnetic Twisting Cytometry (MTC) technique. Ferromagnetic microbeads coated with an integrin specific ligand, (i.e., fibronectin), are used to probe the mechanical (rheological) properties of the cells when a torque is exerted by an externally applied magnetic field [27, 28].

CSK stiffness measurements performed after 30 min (Fig 7A) and 24 h (Fig 7B) of cell exposure to different toxin concentrations reveal a CyaA-dependent increase in CSK stiffness of up to 45% (as compared to that of control cells) after 30 min-exposure to the highest CyaA concentration (10 nM). This increase remains very significant even after 24 h-exposure to CyaA. Noteworthy, the significant decay in CSK stiffness observed after cytochalasin D treatment confirms that the measured stiffness is actin-CSK specific as expected by the use of fibronectin-coated microbeads functionalized for integrins (see Material and Methods). Note that changes in cell mechanical properties measured by MTC which are also known to reflect changes in cytoskeleton internal tension [34] are closely related to cell shape [37, 38].

Cell migration and tissue repair observed during a wound healing test is a hallmark of functional integrity for tissue cells since it involves fundamental cell processes such as adhesion, locomotion, cytoskeleton mechanical properties including intracellular tension, actin polymerization and actomyosin motors activation. We therefore explored whether CyaA intoxication might affect these processes. The wound healing test was carried out on monolayers of A549 alveolar epithelial cells previously exposed for one hour to different concentrations of CyaA (0.5, 5, or 10 nM) and compared to control conditions (no toxin). The cell monolayers were then scratched with a pipette tip of 5 μm diameter to create a wound, and cell migration and repair were followed by microscopy until complete closure of the wounds (the repair area were evaluated every 3 hrs up to 40 hrs). Results presented in Fig 8A reveal that CyaA intoxication drastically slows down both cell migration and overall wound repair. At the low toxin concentration of 0.5 nM, the repair area is significantly reduced at all times tested. At the highest concentration tested (10 nM), the repair is almost totally abolished (> 95%). As expected, the enzymatically inactive CyaA variant CyaAE5 had no effect on the cell migration nor wound repair (Fig 8B). Altogether, our present results indicate that the increase in intracellular cAMP elicited by the CyaA toxin directly impairs the cytoskeletal and mechanical properties of the intoxicated A549 alveolar epithelial cells and reduce their migration and wound healing capacities. As these effects are observed at the low toxin concentrations (0.5 nM) that may be produced in vivo during *B. pertussis* infection [13], our results suggest that the CyaA could also contribute to the local alteration of the epithelial barrier of the respiratory tract, that is an hallmark of *pertussis*.

**Fig 7.**
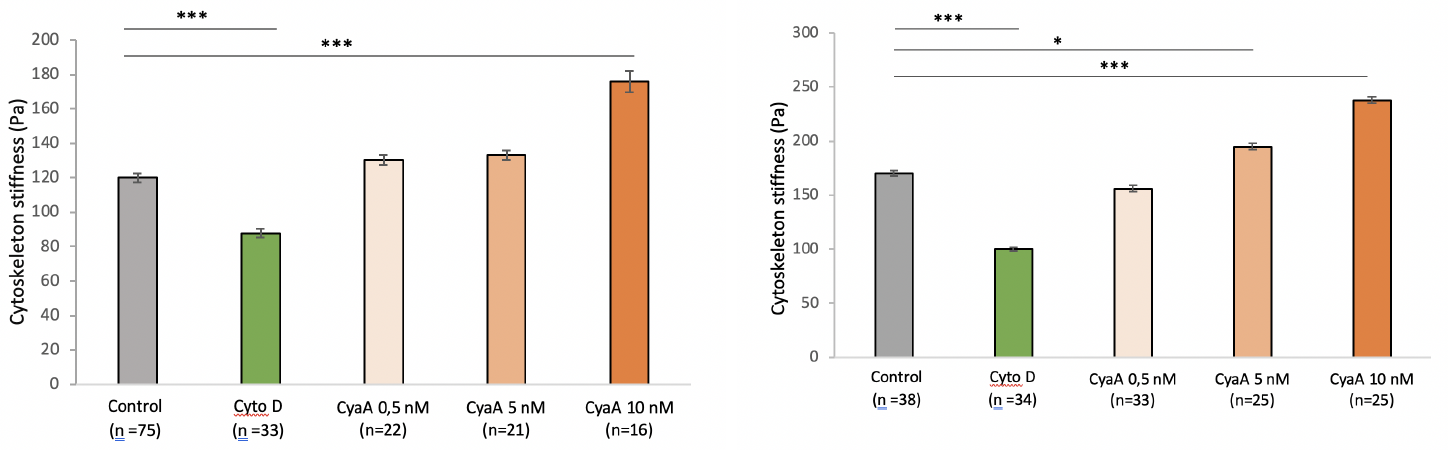
Effect of CyaA on the cytoskeleton stiffness of A549 cells. A549 cells were exposed for 60 min (A) or 24 hrs (B) to the indicated concentrations of CyaA toxin (control=no CyaA) or 10 μM of cytochalasin D (an actin-depolymerizing drug). The CSK stiffness is assessed using the viscoelastic solid-like model (simple Voigt) as described in ‘Material and Methods’. Each MTC measurement is performed on a wide number of cells (7 × 10^4^) probed with an even higher number of beads (about twice). The ‘n’ corresponds to the number of wells. Differences in cytoskeleton stiffness observed between the different groups are quantified by ANOVA test compared to the control conditions. Error bars are ± SEM; * *p* ≤ 0.05; ** *p* ≤ 0.01; *** *p* ≤ 0.001.

**Fig 8.**
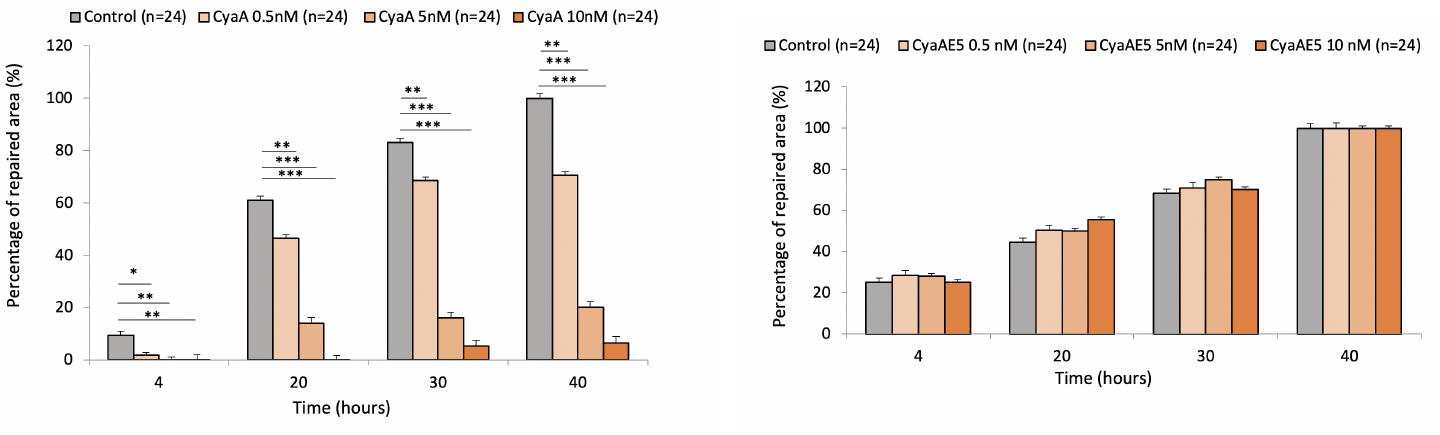
Effect of the CyaA toxin on wound repair of A549 cell monolayers. A549 cells, grown to form a confluent monolayer, were exposed during 1 hr to the indicated concentrations of CyaA toxin (control=no added CyaA) (A) or the enzymatically inactive CyaAE5 protein (B). The cell monolayers were then scratched with a pipette tip (5 μm-distal diameter) to create a wound. Cells are exposed to the indicated concentrations of CyaA or CyaAE5 during 60 min. After washing, cells were further grown in DMEM medium and the repair of the wounds was monitored by microscopy imaging (the wound area was measured with ImageJ) every 3 hrs until complete closure (repaired area = 100%). The data correspond to the percentage of repaired area at different times until complete repair of a wound that occur after 40 hrs in control condition. ‘n’ corresponds to the number of analyzed wells. Error bars are ± SEM; * *p* ≤ 0.05; ** *p* ≤ 0.01; *** *p* ≤ 0.001.

## Discussion and perspectives

While the infection by CyaA of myeloid cells such as macrophages, neutrophils, dendritic cells, natural killer cells, has been extensively studied [4, 39–41], much less is known about the intoxication of non-inflammatory cells lacking the CD11b/CD18 receptor. In these cells, CyaA is able to directly interact with the plasma membrane and translocate its catalytic domain across the lipid bilayer although the precise molecular mechanism of intoxication remains elusive [1, 8, 42, 43].

We previously showed and we confirm here that CyaA can efficiently invade A549 alveolar epithelial cells to trigger a rapid and massive increase in intracellular cAMP. Our present data show that in these cells, CyaA induces a remodeling of actin cytoskeleton, a rapid decrease in size and number of FA sites, resulting in cell rounding and a significant increase in CSK stiffness. These modifications drastically impair the migration capabilities of A549 cells and their ability to repair wounds made in a monolayer of these epithelial cells.

It has already been shown that CyaA invasion results in a deep remodeling of the actinic CSK localized essentially under the plasma membrane [9, 44]. Present data obtained on cellular adhesion as well as previous data obtained on early adhesion in the same cellular system [16] show that CyaA significantly weakens the adhesion system, suggesting that mechanotransduction signaling pathways are affected during the intoxication process.

In our previous study on the early development of adhesion sites in intoxicated cells [16], CyaA was reported to reinforce adhesion by decreasing the strength of newly formed integrin bonds as well as their mutual association through clustering. Thus, alteration of adhesion process and weakening of cell-matrix attachments can be considered as the hallmark of initial intoxication. Here, we show that CyaA intoxication is responsible for a cell rounding process that is likely resulting from the early alteration of adhesion. Such CyaA-induced morphological changes have already been documented in a variety of cultured cell lines [14]. When cells round and cystokeleton is deformed, filamin protein A (an actin-binding protein that crosslinks F-actin) binds to p190RhoGAP and prevents its accumulation in lipid raft. p190RhoGAP is a RhoGAP protein which mediates integrin ligation-dependent inactivation of Rho [45]. Thus, in rounded cells, p190RhoGAP is inactive and hence the Rho activity is presumed to be high [46]. By contrast, in spread cells, p190RhoGAP accumulates in lipid rafts and become active leading to the suppression of Rho activity. Note that Rho’s ability to increase myosin light chain phosphorylation through activation of Rho-associated kinase (ROCK) may feedback to further activate Rho by increasing level of tension within the cytoskeleton [47, 48]. The increase in CSK stifffness presently observed in intoxicated cells is consistent with such an assumption. Activation of Rho GTPases in adhesion signaling involves extensive crosstalk between integrins, Src family kinases, and between individual Rho GTPases themselves [49, 50]. As a matter of fact, changes in cell shape are classically converted into changes in intracellular biochemistry leading to changes in actin-cytoskeleton-dependent control of Rho GTPase activity [51, 52]. Therefore, the balance RhoA and/or Rac1 which is susceptible to affect cytoskeletal structure and cell mechanical properties [53, 54] is likely modified, primarily by the toxin induced change in cAMP levels [55] and secondarily by the CyaA-induced adhesion remodeling which affects cell shape (see adhesion data presently reported in Fig 4 and results obtained in previous studies [14]). By contrast, Kamanova *et al.* [9] reported that in macrophages the increase in cAMP elicited by CyaA causes transient RhoA inactivation that triggers actin CSK rearrangements and phagocytosis inhibition. Incidentally, several toxins deeply modifying cAMP production, (e.g., EDIN of *Staphylococcus aureus*, edema toxin (ET) of *Bacillus anthracis*, or Cholera toxin from *Vibrio cholerae*), induce rupture of the host endothelium barrier and thereby promote bacterial dissemination, by altering the actin cytoskeleton though disruption of actin cables, a phenomena resulting from a change in the balance between F-actin-regulating molecules [56, 57].

It is also recognized that migration, control of morphogenesis, and cell polarity are tightly controlled by Rho GTPases [58]. Present data show that these cellular functions are deeply affected by the toxin. The cytotoxic activity of the toxins caused by actin depolymerisation has already been explained by the role of certain Rho GTPases [59]; Integrin-mediated spreading and focal adhesion maturation develop as a result of a biphasic reaction associated with the relative activities of RhoA and Rac1 [45, 49]. Early adhesion, involving nascent adhesions, is dependent on Rac1 activation and a concomitant suppression of RhoA activity. In contrast, mature focal adhesion and the contractile machinery (stress fibers, internal tension), are reliant on elevated RhoA activity and Rac1 inhibition [60].

Overall, the toxin-induced cellular alterations experimentally observed in our A549 cells are consistent with the assumption of the maintenance of RhoA activity. RhoA is the upstream activator of the serine / threonine kinase RHOK (Rho-kinase) which in cooperation with scaffold protein mDia regulate actin-myosin assembly and contractility as well as actin polymerization resulting in modifications in stress fiber formation and cell motility. We indeed observed on cellular images a persistence of stress fibers up to the highest CyaA concentration, while cytoskeleton stiffness was continuously increased as CyaA concentration was increased. On the other hand, cellular migration was significantly affected after CyaA intoxication in a concentration-dependent manner which suggests that the migratory phenotype which requires Rac1 activation might be limited by intoxication [61]. Moreover, it has been shown that cAMP modulates cell morphology by inhibiting a Rac-dependent signaling pathway [55]. Similarly, the results obtained in our previous study on the early development of adhesion while the same cells were intoxicated with the same toxin [16], have shown a reduction in chemical energy of individual integrin-RGD ligand bonds as well as a diminution by two of integrin association at the level of newly formed adhesion sites (clustering). For these two processes, the role of Rac 1 is fundamental [62] and its potential defect in parallel to the activation of RhoA could be the common cause of these results. Yet, this presumed molecular mechanism remains to be evaluated in future studies.

Another important aspect that may explain the efficient effects of CyaA on the actin cytoskeleton remodeling and mechanical perturbations of the A549 cells is cAMP compartmentalization. Indeed, the specific pathway of CyaA entry, that involves direct translocation of the toxin catalytic domain across the plasma membrane of the intoxicated cells, results in a rapid and preferential production of cAMP in the vicinity of the plasma membrane with a delayed and lower cAMP signal in a perinuclear area [63]. It is well known that microdomains of cAMP have pronounced effects on various signaling processes and major physiological functions in a wide variety of cells [64]. For CyaA, this has been particularly well characterized in T cells where the toxin was shown to efficiently disrupt the immunological synapse and redistribute a number of essential players in the T-cell activation cascade [65, 66]. Remarkably, these effects are not elicited by another bacterial toxin, the anthrax edema factor that produces similar cytosolic levels of cAMP but originating from a perinuclear localization - where the toxin is released from late endosomes [66]. Hasan *et al.* recently reported similar findings on human monocytes [67]. It will be interesting to further delineate the precise role of cAMP compartmentalization in the biological effects of CyaA on the actin cytoskeleton remodeling and mechanical perturbations of epithelial cells evidenced here.

## Conclusion

In summary, we show here that the CyaA toxin is able to efficiently target epithelial cells where the large increase in intracellular cAMP elicits various structural and functional modifications of the cells, including cell rounding, weakening of adhesions, remodeling of actin structures and of cytoskeleton stiffness. This leads to a drastic inhibition of cell migration and wound repair capabilities. These data support the hypothesis that CyaA, in addition to its critical role in blunting the innate immune responses, may also contribute, likely in synergistic manner with other virulence factors, to the local alterations of the epithelial barriers of the respiratory tract, a pathognomonic feature of *B. pertussis* infection.

## Supporting information

**Fig. S1.**
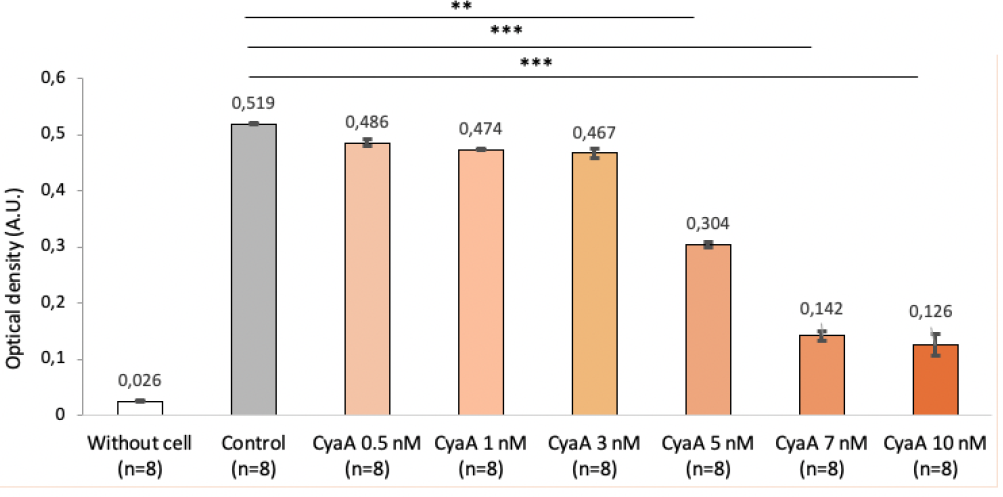
Viability assays of A549 cells exposed to CyaA toxin. A549 cells were grown in monolayer to 90% of confluence and then incubated for 60 min with the indicated concentrations of CyaA. MTT was then added at 0.25 μg/ml and cells were further incubated 4 hours at 37 °C. The medium was removed and replaced by 200 μL of DMSO and the optical density at 550 nm was recorded on a microplate reader. Control corresponds to cells incubated in similar conditions without CyaA. Error bars are ± SEM; * *p* ≤ 0.05; ** *p* ≤ 0.01; *** *p* ≤ 0.001. These data show that the viability of A549 cells is not significantly affected when they are exposed during 1 hr to CyaA concentrations lower than 3 nM, while it is drastically reduced at CyaA concentrations above 5 nM.

**Fig. S2.**
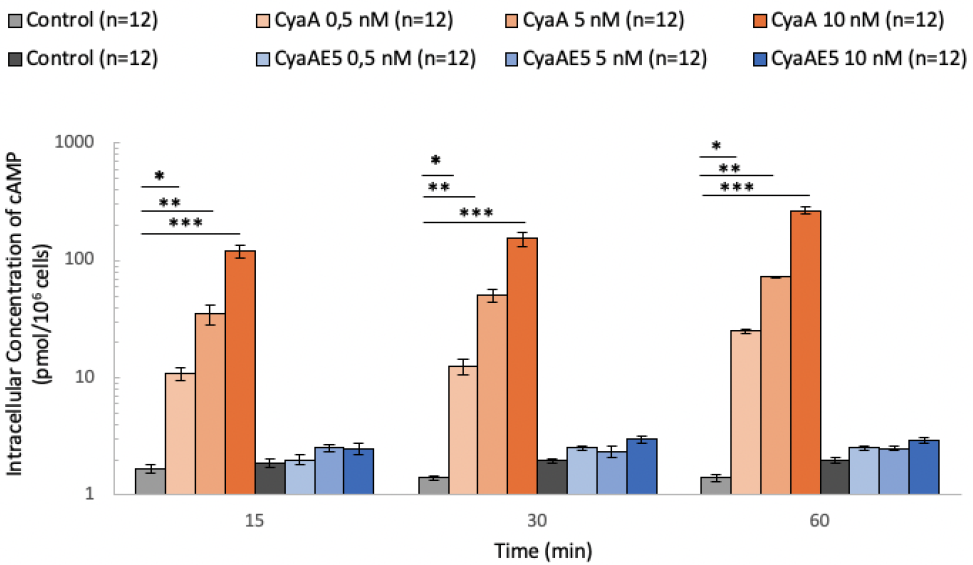
Intracellular cAMP measurements in A549 cells exposed to either CyaA or CyaAE5 toxins. Intracellular cAMP is measured by ELISA assay in A549 cells exposed to CyaA or to CyaAE5, a CyaA variant lacking enzymatic activity, at concentrations 0.5 ; 5 and 10 nM and for 15, 30, and 60 min (n=12 wells). Control conditions correspond to cells incubated without toxin. Error bars are ± SEM ; * *p* ≤ 0.05; ** *p* ≤ 0.01; *** *p* ≤ 0.001. These data show that even the lowest CyaA concentration (0.5 nM) triggers a large increase in intracellular cAMP, that can be observed at the shortest exposure time (15 min) while very high cAMP levels can be reached observed at higher CyaA concentrations. As expected, no significant changes in intracellular cAMP levels are observed when cells are incubated with the enzymatically inactive toxin, CyaAE5.

## Acknowledgments

The authors thank Emilie Bequignon, André Coste, Estelle Escudier, Jean-Francóis Papon, Sofia Andre Dias and Ngoc Minh Nguyen for their helpful advices and fruitful discussion.

